# A genomic panel for studying C3-C4 intermediate photosynthesis in the Brassiceae tribe

**DOI:** 10.1101/2023.01.22.525068

**Authors:** Ricardo Guerreiro, Venkata Suresh Bonthala, Urte Schlüter, Sebastian Triesch, Andreas P.M. Weber, Benjamin Stich

## Abstract

Research on C4 and C3-C4 photosynthesis has attracted significant attention because the understanding of the genetic underpinnings of this trait will support the introduction of its characteristics into commercially relevant crop species. We used a panel of 19 taxa of 18 Brassiceae species with different photosynthesis characteristics (C3 and C3-C4) with the following objectives: (i) create draft genome assemblies and annotations, (ii) quantify the level of orthology using synteny maps between all pairs of taxa, (iii) describe the phylogenetic relatedness across all the species, and (iv) track the evolution of C3-C4 intermediate photosynthesis in the Brassiceae tribe.

Our results indicate that the draft *de novo* genome assemblies are of high quality and cover at least 90% of the gene space. Therewith we more than doubled the sampling depth of genomes of the Brassiceae tribe that comprises commercially important as well as biologically interesting species. The gene annotation generated high-quality gene models, and for most genes extensive upstream sequences are available for all taxa, yielding potential to explore variants in regulatory sequences. The genome-based phylogenetic tree of the Brassiceae contained two main clades and indicated that the C3-C4 intermediate photosynthesis has evolved five times independently. Furthermore, our study provides the first genomic support of the hypothesis that *Diplotaxis muralis* is a natural hybrid of *D. tenuifolia* and *D. viminea*. Altogether, the *de novo* genome assemblies and the annotations reported in this study are a valuable resource for research on the evolution of C3-C4 intermediate photosynthesis.

## INTRODUCTION

Carbon concentrating mechanisms enable plants to reduce photorespiration and improve their photosynthetic efficiency especially under conditions of high temperatures and limited water supply (Sage et al., 2012, Walker et al., 2016, Bellasio and Farquhar, 2019). In C4 photosynthesis, a high CO_2_ atmosphere is achieved in the bundle sheath cells by complex modifications of leaf biochemistry, anatomy and ultrastructure (Hatch 1987). C4 photosynthesis is therefore not only the focus of fundamental research but also crop breeding programs may benefit from a better knowledge of the trait (Schuler et al, 2016). However, our understanding of the genetics underlying C4 photosynthesis is still very fragmented and attempts to introduce C4 traits into agriculturally relevant crop species were only partially successful (Wang et al., 2017, Ermakova et al., 2021). An alternative approach might therefore focus on the understanding of carbon concentration through the glycine shuttle mechanism, a pathway that is supposed to represent an early step during the evolution from C3 to C4 photosynthesis (Rawsthorne et al., 1992; Mallmann et al., 2014). Plants employing the glycine shuttle mechanism are often termed C3-C4 intermediates or C2 species because a C2 compound is exchanged between the cells (Edwards and Ku, 1987, Rawsthorne et al., 1992; Sage et al., 2014). Measurable parameters such as the CO_2_ compensation point or the organelle accumulation in bundle sheath cells usually show intermediate values between C3 and C4 plants (Ku et al., 1991, McKown and Dengler, 2007, Muhaidat et al., 2011). Biochemical, anatomical and ultrastructural modifications in the C3-C4 leaf are therefore likely to be less complex than in C4 plants, easier to understand and, thus, to engineer (Lundgren 2021, Bellasio and Farquhar, 2019).

The photorespiratory cycle describes the recycling of 2-phosphoglycerate (2PG), a toxic metabolite that is formed when Rubisco reacts with oxygen instead of CO2. 2PG is initially converted into glycolate in the plastids and transported into the peroxisome. There it is further metabolized into glyoxylate and aminated into glycine. In the mitochondria two molecules of glycine are converted by coordinated reactions of the glycine decarboxylase complex and the serine hydroxymethyl transferase into one molecule of serine, CO_2_ and NH_3_ (for recent reviews see: Eisenhut et al., 2019, Timm and Hagemann, 2020). Through further reactions taking place in the peroxisome and plastid, serine is deaminated into hydroxypyruvate, then metabolised into glycerate and finally converted into 3-phosphoglycerate, a metabolite that can enter into the Calvin-Benson-Bassham cycle. In C3 species the complete photorespiratory cycle takes place in all photosynthetically active cells of the leaf. Shifting the glycine decarboxylation step exclusively to the bundle sheath cells leads to increased CO2 release in these cells, creating an elevated CO_2_ environment around the bundle sheath Rubisco and considerably reducing the oxygenase reaction in this compartment. The bundle sheath specific localisation of the P-protein from the glycine decarboxylase complex has been shown in C3-C4 species from diverse phylogenetic backgrounds by immunolocalization (Rawsthorne et al., 1988, Schlüter and Weber, 2016, Khoshravesh et al., 2016, Oono et al., 2022). The glycine shuttle biochemistry is accompanied by enhanced centripetal organelle accumulation in the bundle sheath cells (for reviews see: Schlüter and Weber, 2016, Lundgren, 2021). Carbon concentration via the glycine shuttle is less effective than the C4 cycle, but could be advantageous under hot and dry growth conditions, when photorespiration is usually high (Belassio and Farquhar, 2019). Since the C3-C4 related features could represent transitory stages towards C4 photosynthesis, knowledge of their underlying genetic structures could also contribute to the understanding of C4 evolution.

The anatomical and physiological differences between C3 and C3-C4 intermediate species are relatively well studied and characterised in the Brassicaceae genus *Moricandia* (Schlüter et al., 2017). Genetic factors responsible for these differences have mostly been analysed through the lens of transcriptomics (Gowik et al., 2011, Bräutigam et al., 2011; Schlüter et al., 2017, Lauterbach et al., 2017). While transcriptome analysis unravels gene expression patterns, it alone is not sufficient for understanding gene regulatory mechanisms (Conant et al., 2014). Therefore, as an extra layer of information, whole genome assemblies can be used with comparative and quantitative approaches to investigate the regulatory genes and elements, genome duplications and structural variations (Conant et al., 2014, Adwy et al., 20,15, 2019, Schulze et al., 2013). The existence of genome assemblies also facilitates other classical and modern genetic approaches, such as primer design, targeted sequencing, quantitative trait locus (QTL) analysis, pan-genome association mapping and genome editing (Jeong et al., 2019).

Most effort has so far been put into understanding of C4 photosynthesis in phylogenetically disparate species such as maize (Wang et al., 2013, Denton et al., 2017), *Gynandropsis gynandra* (Külahoglu et a., 2014, Reeves et al., 2018) or *Flaveria* sp. (Gowik et al., 2011, Taniguchi et al., 2021) and the implementation of the complete C4 trait into quite distantly related but agriculturally relevant C3 species such as rice (von Caemmerer, 2012; Schuler et al., 2016, Ermakova et al., 2021). Understanding how to convert C3 into C3-C4 photosynthesis is less challenging and could already produce commercially relevant yield gains (Schuler et al., 2016; Weber & Bar-Even, 2019, Lundgren, 2021). In this context, the Brassicaceae family is intriguing as it contains the genetically very well characterized model species *A. thaliana* and commercially relevant species such as *B. napus* (canola) and *B. oleraceae* (cabbage). In addition, this family also includes multiple C3-C4 intermediate evolutionary lineages (Apel et al., 1997; Sage et al., 2011). Hence, Brassicaceae species are ideal for investigating C3-C4 evolution in a pan-genomic context to understand the differences in gene regulation and studying convergent evolution. In addition, Brassicaceae species are known to produce fertile progenies in interspecific crosses (Kaneko & Bang, 2014; Ueno et al., 2003). Such progenies can be be helpful for unravelling the inheritance of C3-C4 intermediacy and for transferring the genes of interest to relevant crops.

The main aim of this study is to establish the genomic resources that enable comparative genetic and genomic research on C3-C4 intermediate photosynthesis. In detail, the objectives of our study were to:

1. create draft genome assemblies and annotations of 19 closely related Brassiceae taxa with different photosynthesis characteristics (C3 and C3-C4),
2. quantify the level of orthology using synteny maps between all pairs of taxa,
3. describe the phylogenetic relatedness across all the taxa, and
4. track the evolution of C3-C4 intermediate photosynthesis in the Brassicaceae family.

## MATERIALS AND METHODS

### Genetic material

We collected seeds for 18 Brassiceae species (19 taxa) from various gene banks (Suppl. Table 1), for which the genome sequences were unavailable. Thereby we considerably increased the coverage of this tribe in which C3-C4 intermediacy has been reported previously. A subset of the above mentioned taxa was selfed one to several times to reduce heterozygosity and facilitate genome assembly later.

### Linked-read library preparation and sequencing

For all 19 taxa, linked-read sequencing was performed using either 10x (Zheng et al., 2016) or stLFR (Wang, O et al, 2019) technologies (Suppl. Table 1). Initially, for 15 taxa (Suppl. Table 1), DNA was extracted with the DNeasy Plant Mini Kit (QIAGEN) following the manufacturer’s instructions and size-selected for fragments larger than 40 Kbp using BluePippin (SAGE Sciences). Quality control of the size-selected DNA was performed on Qubit and TapeStation. A 10x linked-read library (Zheng et al., 2016) was created for each taxa using 1 ng of DNA as recommended by the manufacturer. Sequencing was performed on the HiSeq3000 sequencer with pair-end mode by Novogene.

For the remaining four taxa (Suppl. Table 1), stLFR linked-read libraries (Wang et al., 2019) were prepared by BGI from tissue samples using MGIEasy stLFR Library Prep Kit (MGI, Shenzhen, China). The libraries were sequenced on BGISEQ-500 (100 bp and pair-end) by BGI. In addition, we re-sequenced one species due to unsatisfactory quality of 10x data, using stLFR link-read technology as mentioned above (Suppl. Table 2).

### Long-read library preparation and sequencing

Complementary long-read data was generated for a subset of seven taxa (Suppl. Table 1) to improve the *de novo* genome assemblies. PacBio SMRTbell libraries were prepared as recommended by Pacific Biosciences (SMRTbell Template Prep Kit 1.0 <u>SPv3</u>), including a size selection on Blue Pippin to remove fragments lower than 10 Kb. Sequencing was performed on Sequel with 2.0 Binding Kit and sequencing chemistry for 10 h, or 3.0 Binding Kit and sequencing chemistry for 20h, as recommended by Pacific Biosciences. Oxford Nanopore libraries (Suppl. Table 1) were prepared from purified high molecular weight DNA extracted from leaf tissue by precipitation of DNA-CTAB complexes (Arseneau et al., 2017; Xin & Chen, 2012). In a second step, CTAB was removed with ethanol, and the co-purified RNA was digested by RNAse treatment. Afterwards, the DNA was again purified by binding it to AMpure PB (Pacific Biosciences) beads, washing the beads in ethanol and then resolving the DNA. Sequencing was performed by GridION and PromethION flow cells by GTL Düsseldorf.

### Estimation of genome size, heterozygosity and repeat content

From the linked-read libraries of each taxa, the 21-mers were extracted using Jellyfish (version 2.1.3) (Marçais & Kingsford, 2011). Genomescope (www.genomescope.com) was then used to estimate genome size, heterozygosity and repeat content, setting the maximal kmer coverage parameter to 10,000.

### Genome assembly

The supernova assembler v2.1.1 (Weisenfeld et al., 2017) was used to assemble both 10x and stLFR linked read data to pseudohaploid assemblies. Long Ranger v2.2.2 (Ott et al., 2018) was used with default parameter settings to map the linked reads to respective *de novo* genome assemblies. Purge Haplotigs v1.1.0 (Roach et al., 2018) was used to reduce the under-collapsed haplotigs in all *de novo* genome assemblies. Deduplication was not possible for *D. muralis* due to Long Ranger failing to map the linked-reads. BUSCO v3.1.0 (Simão et al., 2015) was used to estimate based on the eudicot_db10 database the completeness of the *de novo* genome assemblies before and after reducing the under-collapsed haplotigs.

PacBio long reads for *E. sativa* and *D. erucoides* were assembled with Canu v1.8 (Koren et al., 2017) using default parameters except for corOutcoverage=200 and correctedErrorRate=0.15 and discarding reads shorter than 1,000 bp. To deal with the higher error frequency of long reads, the Canu assemblies were polished using Pilon v1.22 (Walker et al., 2014) with the less error-prone linked reads by mapping in two iterations. The polished and purged PacBio assemblies were further scaffolded with the LINKS v1.8.7 – ARCS v1.1.1 pipeline (Warren et al., 2015; Yeo et al., 2018) that performs misassembly correction with Tigmint v1.1.2 using linked reads (Jackman et al., 2018).

Oxford Nanopore long reads data obtained for *D. acris, D. harra, H. incana* HIR3, *M. sinaica* and *M. spinosa* were basecalled with Guppy v5.0.11 (Wick et al., 2019). The resulting reads were then trimmed on the first 50 bp and filtered with NanoFilt v2.6.0 (De Coster et al., 2018) on a minimum length of 1000 bp and minimum average phred-64 quality score of 10. The high-quality reads were subsequently used for scaffolding stLFR assemblies with LINKS v1.8.7 (Warren et al., 2015).

Assemblies for *Gynandropsis gynandra* (https://genomevolution.org: id58728), *Moricandia arvensis* and *M. moricandioides* (Lin et al., 2021) were obtained directly from collaborators. The *Moricandia* assemblies were also polished with the 10x reads from our study. Additional assemblies were available from NCBI (Suppl. Table 2).

Assembly statistics such as N50-90, L50-90, assembly size and contig number were calculated with a custom python script for each finalized genome assembly.

### Ploidy estimation

We used nQuire (retrieved in December 2022) (Weiss et al., 2018) to estimate the ploidy of our genomes by analyzing the frequency distribution of biallelic variant sites of reads mapping to BUSCO genes. We generated a histogram of read mapping depths and applied nQuire’s denoise tool, which uses a GMMU approach to remove a uniform baseline from the histogram. The variations in read mapping depth were then used with a Gaussian Mixture Model to generate a log-likelihood under diploid, triploid, tetraploid and free models.The smallest of the delta-log-likelihoods between the free model and the fixed models was taken as the most likely ploidy.

### Transcriptome Assemblies

RNA-Seq data for *E. sativa* (SRR6454139), *H. incana* (SRR11638396), *D. tenuifolia (*PRJNA904765) and *D. viminea* (PRJNA904804) were downloaded from the Sequence Read Archive (SRA) database at NCBI, while for *M. arvensis* and *M. moricandioides* were obtained from Schlüter et al. (2017). Trimmomatic v0.39 (Bolger et al., 2014) was used to trim adapters and low-quality reads. Additionally, reads shorter than 36 bps were discarded. The high-quality RNA-Seq reads were then assembled with Trinity v2.11.0 (Haas et al., 2013).

### Repeat annotation

We performed *de novo* repeat identification using Marker-P guidelines (Campbell et al., 2014). Briefly, Mite-hunter (Han & Wessler, 2010) with the default parameters were used to identify Miniature inverted-repeat TEs (MITEs) and LTRharvest v1.5.9 (Ellinghaus et al., 2008) with the default parameters for *de novo* predictions of LTR (Long Terminal Repeat) retrotransposons. Finally, RepeatModeller v1.0.11 (Smit & Hubley, 2015) with default parameters was used to build a *de novo* repeat library and RepeatMasker v4.0.9 (Smit et al., 2015) was used to mask identified repeats in respective genome assemblies. In addition, repeat annotation was also performed for 13 publicly available species (Suppl. Table 2).

### Gene structural annotation

We used protein sequences of all Brassicaceae species available from the UniProt database (The UniProt Consortium, 2021), excluding protein sequences with low evidence levels (Uncertain and Predicted). We searched the UniProt database on 02/10/2020 using the following parameters: taxonomy: Brassicaceae NOT existence: “Uncertain [5]” NOT existence: “Predicted [4]” OR reviewed: yes). We reduced the sequence identity to 95% between any protein sequences present in the downloaded dataset using CD-HIT (Li & Godzik, 2006; Fu et al., 2012), and the resulting 114,295 protein sequences were used in following gene structural annotation.

Gene structural annotation was performed with Maker2 (Campbell et al., 2014) in two steps. First, potential genes were annotated based on alignments with the protein sequences in our protein database and the transcript sequences assembled for individual species. Second, the annotated genes were fed to SNAP (Korf, 2004) and Augustus (Hoff & Stanke, 2019) to predict gene structure across all taxa. The model training was performed with Nextflow-abInitio, made available by National Bioinformatics Infrastructure Sweden (NBIS), and the trained models were provided in the second run of Maker2.

We initially annotated six taxa: *D. tenuifolia* (C3-C4), *D. viminea* (C3), *H. incana* HIR1 (C3-C4), *M. arvensis* (C3-C4) and *M. moricandioides* (C3) using publicly available RNA transcripts (Mabry et al., 2020; Schlüter et al., 2017) in addition to the protein database mentioned above. The resulting predicted proteins were filtered for AED values smaller than 0.5 and a length > 49 amino acids (the minor 1st-percentile protein lengths in *Arabidopsis*). The resulting protein sequences were added to the above-created protein sequence dataset, followed by reducing the sequence identity to 95% using CD-HIT (Li & Godzik, 2006; Fu et al., 2012). This final protein sequence dataset contained a total of 284,999 proteins. It was used as the only evidence to systematically annotate all genomes of this study with Maker2, including species with existing annotations and the six species we initially annotated. This systematic annotation was done to avoid bias in the downstream analyses with different annotation qualities (Trachana et al., 2011).

### Gene functional annotation

Functional annotation for the predicted proteins was performed using the Automated Assignment of Human Readable Descriptions (AHRD) (https://github.com/groupschoof/AHRD). The AHRD pipeline assigns gene descriptions, Pfam domains (El-Gebali et al., 2019) and Gene Ontology (GO) annotations (Lewis, 2005; Barrel, 2009) for each gene based on InterPro Scan (Zdobnov & Apweiler, 2001) and BLASTp (Altschul et al., 1990) searches. The BLASTp searches were performed against the Araport11 (Cheng et al., 2017), Swiss-Prot (UniProt Consortium, 2021) and trembl_plants (O’Donovan, C. et al., 2002) databases (downloaded in 02/2019).

Transposable element (TE) related genes were identified and excluded from the gene annotation (cf. Jayakodi et al., 2020). A predicted protein was labeled as TE-related and filtered out of the protein sequences if at least two out of three fields of the AHRD output, such as AHRD descriptions, the best-blast-hit description or Pfam annotation, were associated with transposable elements. A list of the terms used for filtering is provided in Suppl. Table 3.

### Orthology map and species tree

The TE-filtered protein sequences were analysed by Orthofinder v2.5.1 (Emms and Kelly, 2015, 2019) for orthology identification. Multiple sequence alignments for identified hierarchical orthogroups (HOGs) were produced with MAFFT v7.471 and used for creating gene trees with RAxML v8.2.10 with the PROTGAMMALG substitution model. The gene trees of all HOGs were fed to ASTRAL-pro with default parameters (Zhang et al., 2020) for generating a multi-species coalescent-approach-based species tree.

### Synteny analysis

A pairwise homology search was performed using BLASTp (Altschul et al., 1990) followed by predicting synteny of genes across all taxa using MCScanX (Wang et al., 2012). A heatmap was generated to visualise the percentage of syntenic genes conserved across all taxa. Finally, we assessed whether the assembly quality has confounding effects on synteny between a pair of taxa by computing Pearson correlations (Suppl. Figures 7 and 8).

## RESULTS

### Genome size estimation

We estimated the genome size for all taxa included in this study using a k-mer based approach. The genome size estimation failed for *D. acris* (Suppl. Table 4). The highest genome size estimation was observed for *D. muralis* and the highest heterozygosity level for *M. spinosa*. Our genome size estimates were 50.8% to 85.3% smaller than reported in the literature, with the largest deviations observed for *B. tourneforti* and *H. incana* HIR3.

### *De novo* genome assemblies

This study reported 19 *de novo* genome assemblies for 19 taxa of 18 Brassiceae species (Table 1). Seven of the 19 assemblies originated from combining long- and linked-read data, while the other 12 were pure linked-read-based assemblies (Suppl. Table 1). The assembly size ranged from 271.11 Mbp for *B. tourneforti* to 884.18 Mbp for *D. acris*. The number of scaffolds ranged from 839 for *D. muralis* to 62,651 for *D. harra*. The assembly quality measurements, such as L50 values, ranged from 7,247 scaffolds for *B. gravinae* to 7 for *B. tournefortii*, while N50 values ranged from 16.5 Kbp for *B. gravinae* to 14.4 Mbp for *B. tourneforti* (Table 1). The genome completeness, assessed by BUSCO against the eudicot_db10 database, ranged from 91% for *B. gravinae* to 99% for *D. muralis*. Except for the allotetraploid *D. muralis*, all assemblies had duplication levels between 12 and 28.1% (Figure 1, Suppl. Table 5).

**Table 1.**
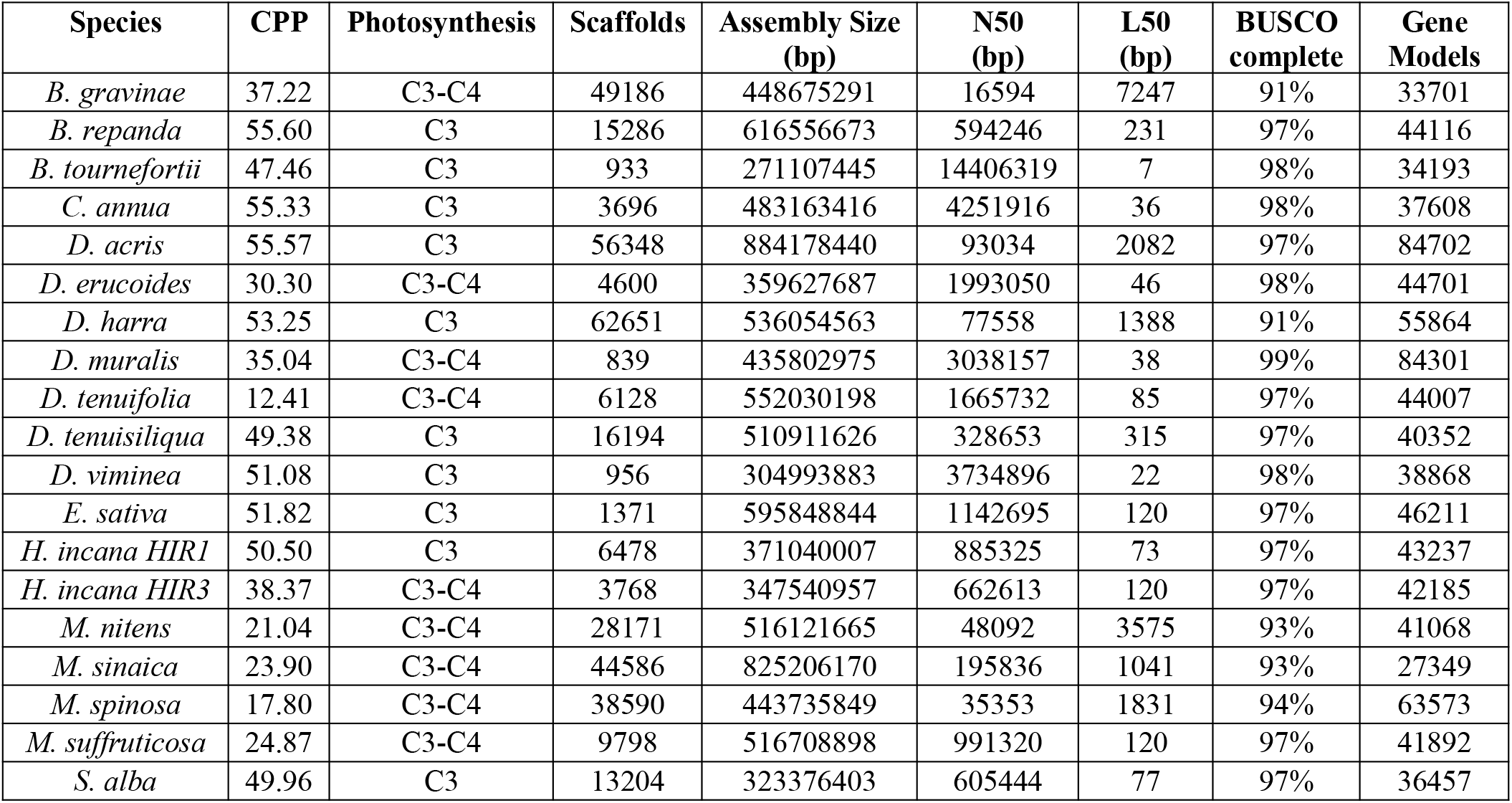
Summary statistics of genome assemblies developed in our study along with CO2 compensation point (CPP) and inferred photosynthesis type from Schlüter et al. (Submited).

**Figure 1:**
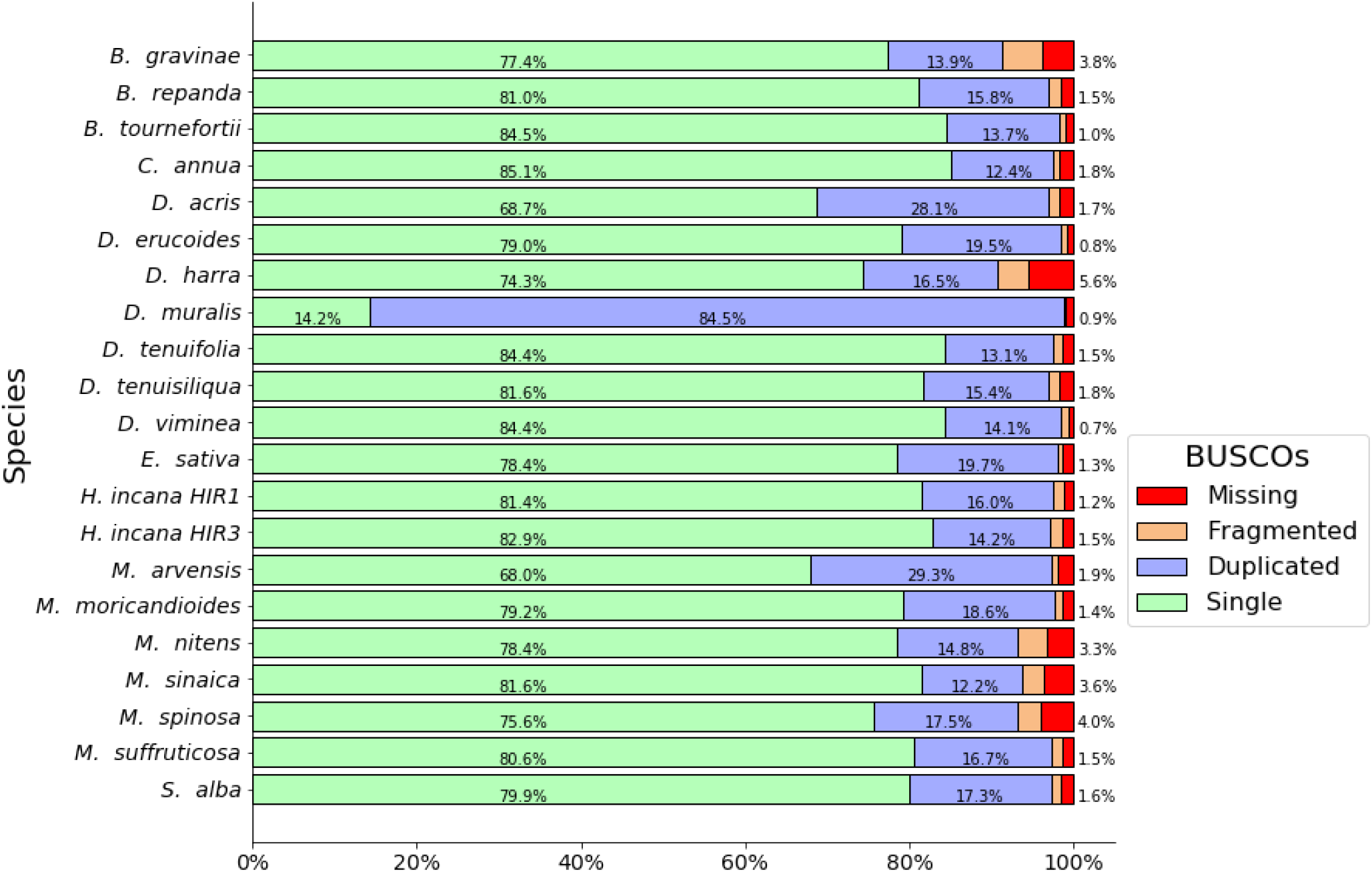
Assessment of the completeness of the 19 *de novo* genome assemblies using BUSCO with eudicot_10db database.

In addition, for downstream analyses, the 19 assemblies generated in this study for taxa from the Brassiceae tribe were complemented with 13 publicly available genome assemblies originating from the Brassicaceae family (Suppl. Table 2). Similarly to our *de novo* assemblies, the fragmentation level and gene completenessvaried across the 13 literature assemblies (Suppl. Table 6).

### Ploidy estimation

The ploidy of the 19 new assemblies was estimated with nQuire based on the frequency distribution of biallelic variant sites of reads that mapped to BUSCO genes. The resulting estimations were diploid for all assemblies except for *D. muralis* and *M. spinosa*, for which the estimations were tetraploid, and for *D. harra, B*.*tournefortii and D. viminea*, where the results were unclear (Suppl. Figure 1).

### Repeat annotation

Repeat content analysis identified an average of 1,937 unique interspersed repeat families in all genome assemblies, including the ones publicly available. The number of notable repeat families ranged from 171 for *R. raphinastrum* to 2810 for *M. arvensis*. Further, an average of 43% of the respective genome assemblies were masked for gene annotation. The lowest proportion of genome assembly was masked for *R. raphinastrum* (6.35%), while the highest amount of genome assembly was masked for *G. gynandra* (65.22%) (Suppl. Table 7).

### Gene annotation

The *de novo* gene annotation was performed for all 19 genome assemblies from this study as well as the publicly available genome assemblies of 13 species using the same method to facilitate comparisons across taxa. The annotations produced a median of 45,408 gene models per assembly, ranging from 22,318 gene models for *A. thaliana* to 113,686 gene models for *B. napus* (Suppl. Table 8) with an average Annotation Edit Distance (AED) score of 0.186. The annotations with the lowest cumulative AED scores were that of *D. acris* and *M. spinosa* (Figure 2). In contrast, the annotations with the highest cumulative AED score were that of *M. moricandioides, M. sinaica* and *S. alba* (Figure 2). An average of 3,648 gene models per assembly were discarded due to high AED scores (>0.5) or their small size (< 50 amino acids). In addition, an average of 1,937 gene models per assembly were discarded due to their functional annotations related to transposable elements (TE). The final gene models for each assembly retained an average of 90.9% BUSCO genes, except *for B. gravinae* (65%) (Suppl. Figure 2). In addition, gene length distribution analysis revealed that the mean and median gene lengths were 1,880 bp and 1,441 bp across taxa, respectively (Suppl. Figure 3). In contrast, the bigger difference between the mean and median inter-genic distances (6,662 bp vs. 2,176 bp) compared to the gene length suggested a more skewed distribution of the former (Suppl. Figure 4).

**Figure 2:**
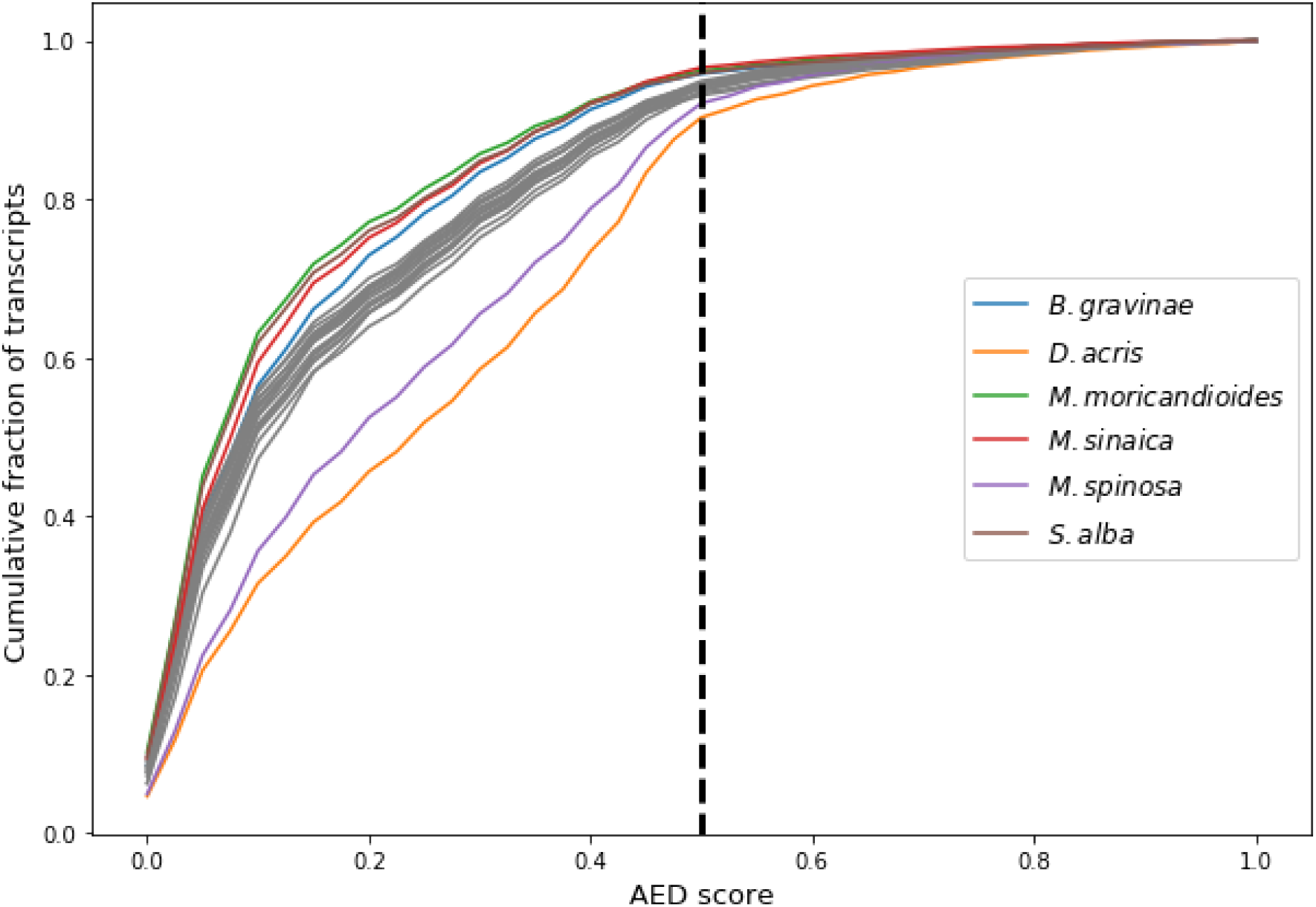
Cumulative annotation edit distance (AED) score of gene annotations for the 19 *de novo* assemblies and 13 publicly available genome assemblies. Highlighted in colour are the assemblies with the highest and lowest proportion of quality gene annotations (AED <0.5).

The length of the upstream sequences for all genes were measured in each assembly by measuring the sequence length from the transcription start site until interruption due to contig end or insertion of Ns. We found that the assemblies of *B. gravinae, D. harra* and *M. sinaica* contained a high percentage of genes with short upstream sequences (<1Kbp). In contrast, all other assemblies were characterized thereby that for the majority of genes very long (>30Kbs) upstream sequences were available (Suppl. Figure 5).

### Orthology and synteny map

Orthofinder clustered about 98% of all protein sequences (1,372,043) from all 32 assemblies into 42,928 orthogroups (HOGs) (Suppl. Table 9). Each HOG indicates a set of homologous genes descended from all taxa’s last common ancestor gene (Emms & Kelly, 2019). Of the 42,928 HOGs, 22,694 were present in less than ten assemblies and were filtered out in order to avoid potential biases when creating the phylogenetic tree. After this filtering step, most assemblies had a median of one gene per HOG (Figure 3A). Exceptions with a higher median of two or three were observed for the tetraploid species and some diploid species (Figure 3A), all of which contained a higher number of total genes (Figure 3B). The percentage of single-copy genes varied from 5.9% in *A. thaliana* to 60.1% in *B. napus* (Figure 3B). Most commonly, HOGs existed in a) all assemblies; b) all but one assembly; c) *D. muralis* and *D. tenuifolia* or *D. viminea*; d) *B. napus* and *B. oleraceae* (Suppl. Figure 6).

**Figure 3:**
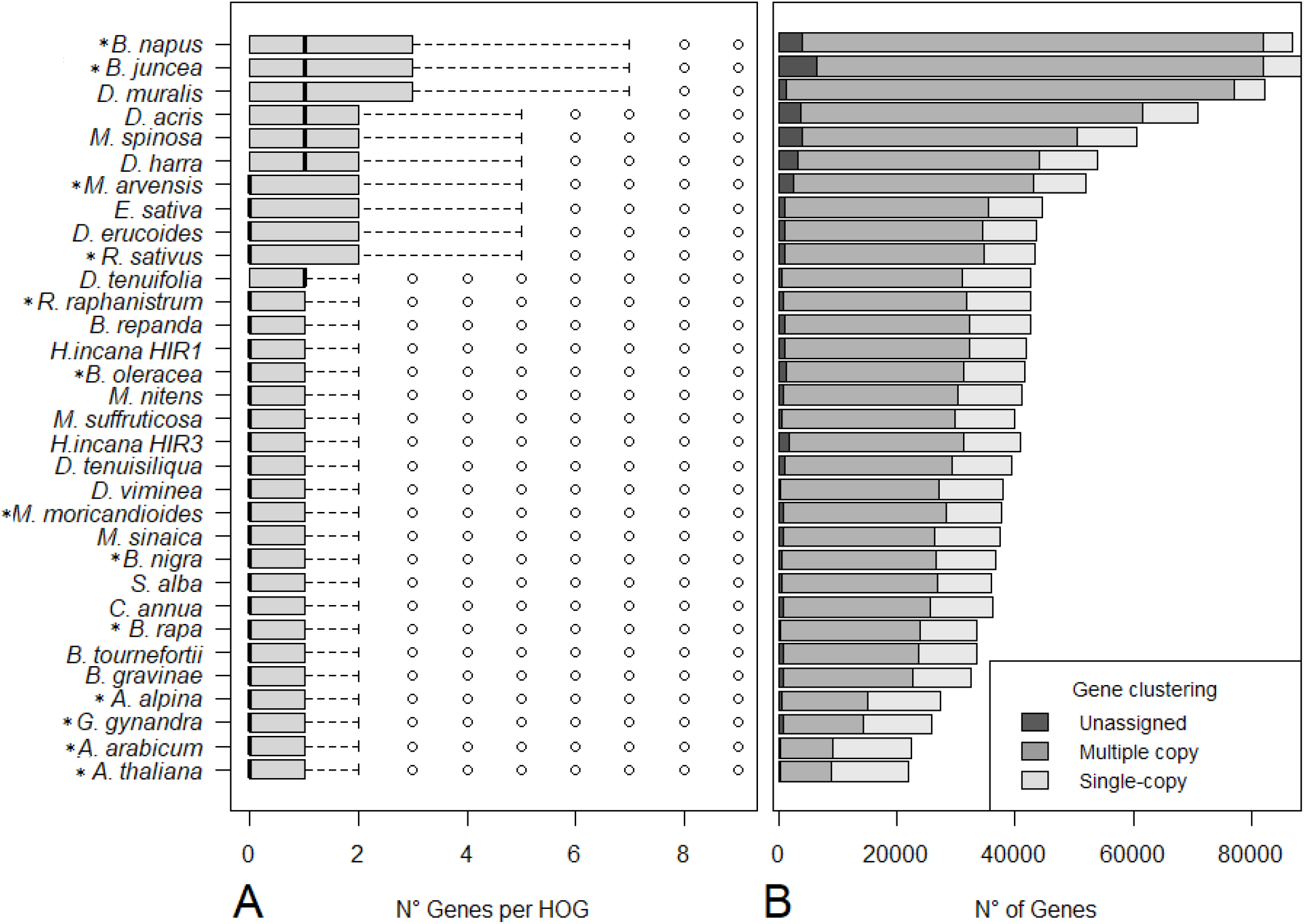
A) Distribution of gene numbers per hierarchical orthogroups (HOG) per taxa in all 32 assemblies B) the total number of unassigned genes, assigned together with other copies or as a single copy to an orthogroup. Species with previously available genome assemblies are marked with an asterisk (*).

We created a synteny map for all 32 assemblies by computing pairwise collinear genes and observed a high conservation of syntenic genes between most taxa. Of particular interest was the high extent of synteny of *D. muralis* with *D. tenuifolia* as well as *D. viminea*. In contrast, *B. gravinae, D. acris, D. harra, M. sinaica* and *R. raphanistrum* showed a very low syntenyagainst all other taxa (Figure 4). Therefore, we performed a correlation analysis to quantify the influence of assembly quality measured as the number of scaffolds (i.e., assembly fragmentation) on the conservation of synteny between each pair of taxa (Suppl. Figure 7). A negative correlation (−0.503 or -0.739 for the number or percentage of collinear genes) between assembly quality (fragmentation) and synteny was observed, indicating that the above reported low synteny can be explained by differences in assembly quality.

**Figure 4:**
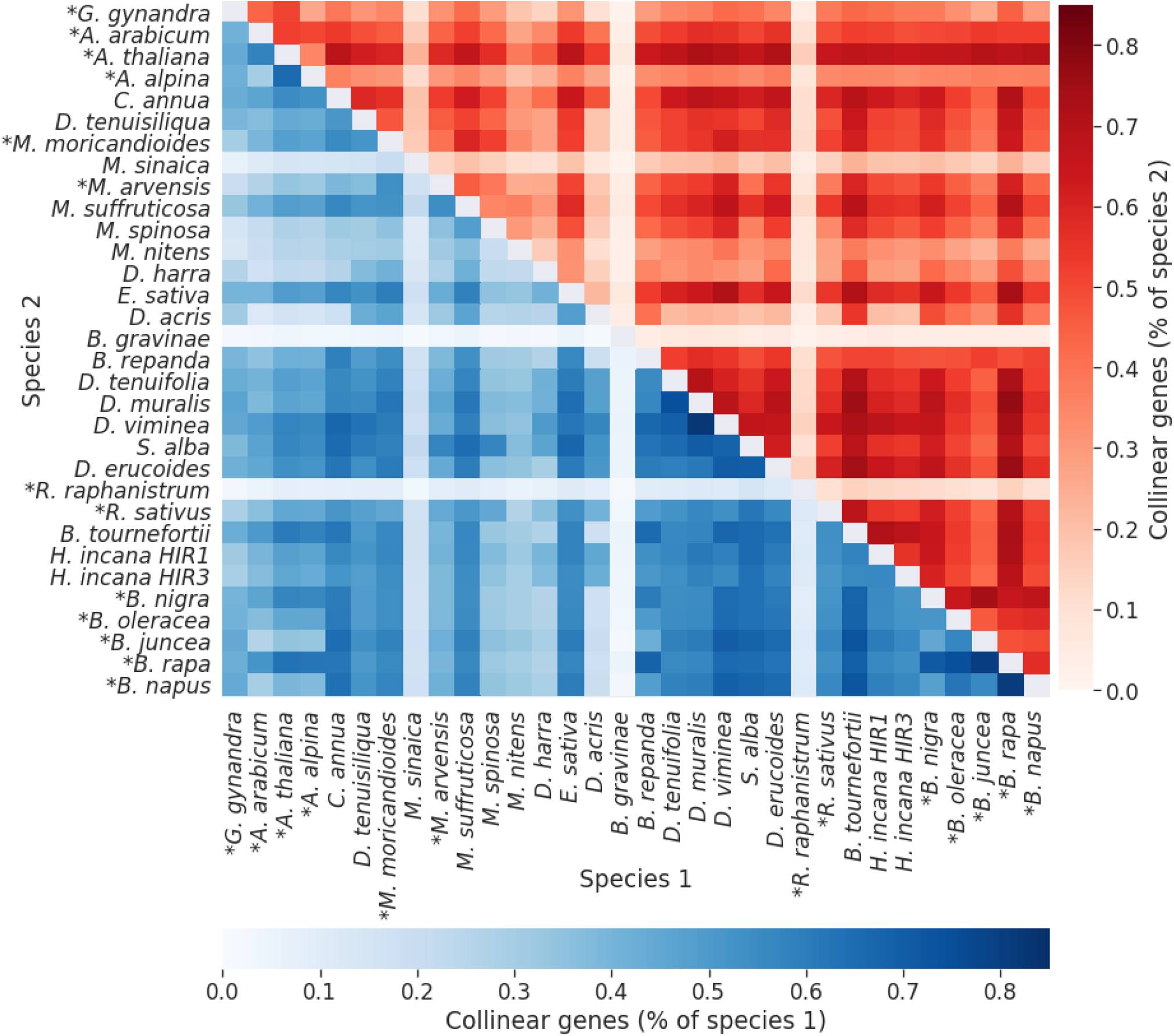
Heatmap of the percentage of syntenic genes between each pair of species with species 1 as reference (below the diagonal) and species 2 as reference (above the diagonal). The species were sorted according to their position in the phylogenetic tree. Species with previously available genome assemblies are marked with an asterisk (*).

### Phylogenetic tree

We established a genome-wide phylogenetic tree for all species included in this study by using ASTRAL-pro with 27793 HOGs. The phylogenetic tree contained two main clades (Figure 5): one clade comprising the taxa of the *Moricandia* genus and most taxa of the *Diplotaxis* genus, as well as *E. sativa*. In contrast, the other clade contained several *Brassica* taxa and the *Raphanus* and *Hirschfeldia* genera. As outgroup, we used *G. gynandra* from the *Cleomaceae* family, which diverged from the Brassicaceae ∼40 million years ago (Edger et al., 2015). The *Moricandia* genus was monophyletic, while the *Diplotaxis* and *Brassica* genera species were dispersed across the phylogenetic tree (Figure 5). When integrating the information of the CO2 compensation points from Schlüter et al. (submitted) in the phylogenetic tree, the result indicated that the C3-C4 intermediate photosynthesis might have developed five times independently (Figure 5).

**Figure 5:**
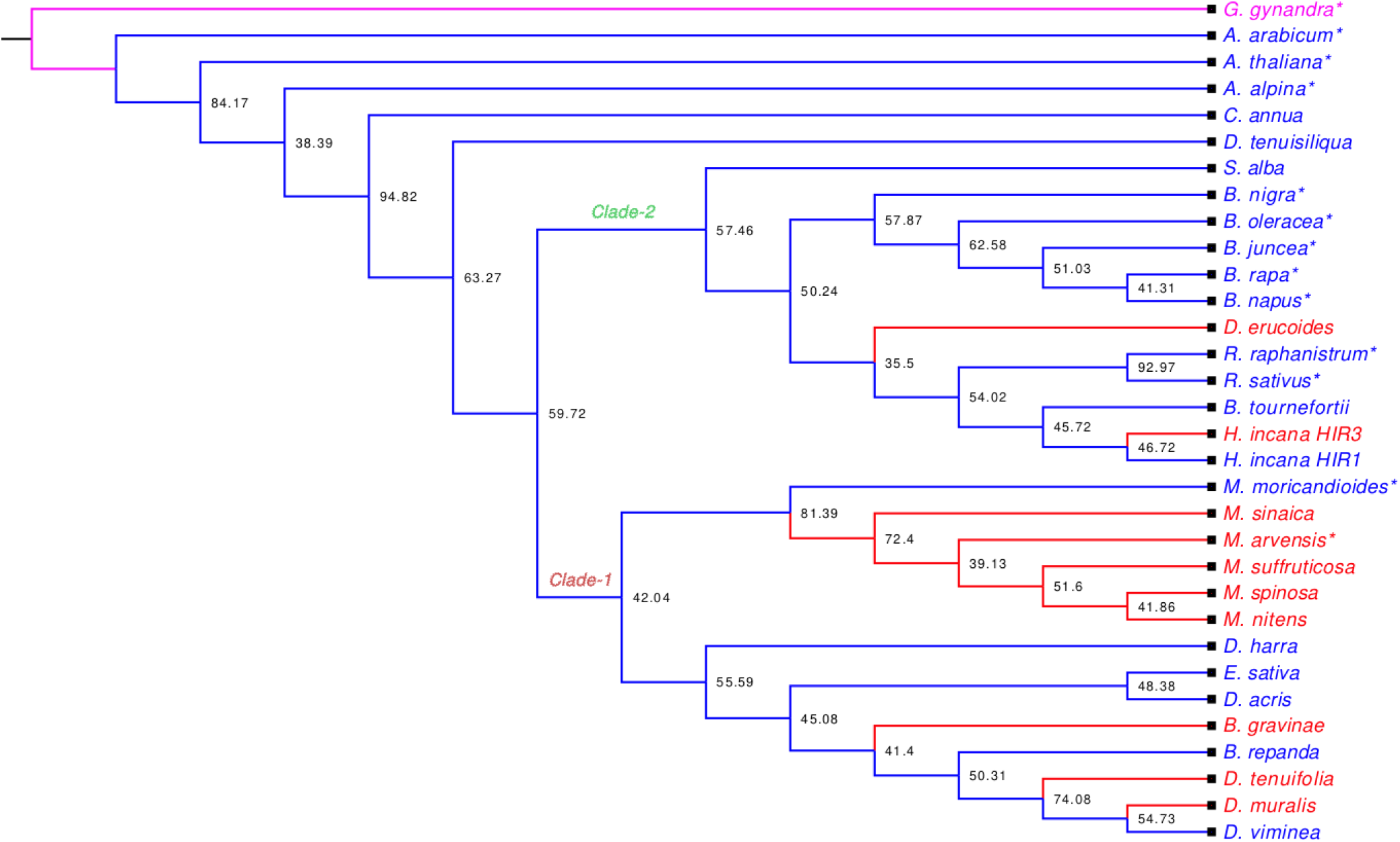
Species tree created using a multi-species coalescent-based approach with *G. gynandra* (C4 photosynthesis) as an outgroup species. The colour code indicates the photosynthesis type inferred from CPP values (Schlüter et al., submitted). Blue colour indicates taxa with C3 photosynthesis, while red colour indicates C3-C4 photosynthesis. Species with previously available genome assemblies are marked with an asterisk (*).

## DISCUSSION

The main aim of this study was to establish the resources that enable genomic comparisons and investigations of the evolution of C3-C4 intermediate photosynthesis within the Brassicaceae family and especially the Brassiceae tribe. For such analyses, not only dense sampling of C3-C4 intermediate species but also of closely related C3 species is required in order to be able to separate signal from noise. Therefore, we have included in this study all species of the Brassiceae tribe which genome was not yet sequenced and for which we were able to obtain seeds.

### High-quality draft *de novo* genome assemblies and annotations

Overall, the genome assemblies generated in this study were fragmented, with varying levels of contiguity and quality. Nevertheless, with at least 91% complete genes identified by BUSCO our assemblies captured most of the gene space (Figure 1), which indicates suitability for comparative genomic analysis. The duplication rates are relatively low, except for *D. muralis* at 84% (discussed below) and *D. acris* at 28% (Figure 1). No k-mer estimation of heterozygosity was possible for *D. acris (Suppl. Table 4)*. Furthermore, all presented assemblies meet the minimum requirement of an N50 larger than average gene length (Yandell & Ence, 2012). Even the N90 values of all assemblies were higher than the average gene length of around 2000 bps (Suppl. Figure 3 and 4). Therefore, the quality of the genome assemblies in this study is comparable or higher than the assemblies available for other Brassicaceae species with similar genome sizes (e.g., Haudry et al., 2013; Lin et al., 2021; Moghe et al., 2014).

The most contiguous assembly was realized in our study for *D. tenuisiliqua* (Suppl. Tables 1 and 4). This is presumably due to the two generations of selfing performed prior to sequencing (cf. Li & Harkess, 2018). In addition, the assembly of *D. muralis* reached satisfactory contiguity after two generations of selfing. Coincidentally, an assembly for *H. incana (Nijmegen)* with six generations of selfing has just been released (Garassino et al., 2022) with an N50 value of 5.1 Mb, which is considerably longer compared to our assemblies with N50s of 885 Kb and 663 Kb for *H. incana* HIR1 and *H. incana* HIR*3* accessions, respectively, that were sequenced without selfing them before.

The assemblies developed in our study result mostly from single linked-read libraries. Contiguity and completeness can be further improved by scaffolding and gap-filling using low-coverage long-read sequencing data, as we illustrated with *H. incana* HIR1, *D. acris, D. harra* and *M. sinaica*. In contrast, we improved the long-read assemblies by scaffolding and polishing with linked-read data for *E. sativa* and *D. erucoides*.

Although the assemblies created in this study are scaffold-level, the assessment of the lengths of the upstream sequences of the transcription start sites (TSS) of annotated genes showed that for a high proportion of annotated genes upstream sequences are available for most of the species (Suppl. Figure 5). Hence, the dataset generated in this study enables analyses downstream from transcriptomics studies, such as the deep analysis of cis-regulatory motifs. Such an analysis could prove crucial in advancing the knowledge about differences underlying gene regulation mechanisms between C3 and C3-C4 intermediate species.

Despite the lack of transcriptional data for all 19 taxa, our *de novo* gene annotation strategy resulted in an average of 46,546 gene models with high quality (AED <= 0.5) across taxa (Suppl. Table 8). The number of gene models for each taxa was comparable to the number of gene models in the gene annotation of the publicly available 13 species that complemented our study.

### Conservation of genes across Brassicaceae species

Resolving the orthologous relationships between species is fundamental to comparative genomics.. We grouped around 98% of annotated genes into orthogroups (Suppl. Table 9), where 65.18% were grouped into 11072 orthogroups present in at least 31 of the used 32 taxa (Suppl. Figure 6). Notably, some orthogroups were absent in species due to low-quality assembly, as evident with *B. gravinae* (Suppl. Fig. 6, Suppl. Table 5), or large phylogenetic distances, as evident from *G. gynandra* and potentially with *A. arabicum* (Figure 5). In contrast, some orthogroups were present exclusively in a specific taxa (Suppl. Fig. 6,Suppl. Table 9). Moreover, these species-specific orthogroups often contained transposable element-related genes (Jayakodi et al., 2020).

To illustrate the possibility of comparing gene regions between pairs of species, we generated a synteny map. As previously mentioned, we consider the particularly high synteny values for *Diplotaxis muralis, D. tenuifolia* and *D. viminea* (Figure 4) as evidence of its hybrid origin. Additional high conservation of synteny is visible across many taxa, with exceptions for *B. gravinae, D. acris, D. harra, M. sinaica* and *R. raphanistrum* (Figure 4). These species had lower assembly contiguity, which correlated strongly with synteny of genes between species (Suppl. Figure 7). When disconsidering these species, the Pearson correlation coefficient between synteny colinear gene number decreased from -0.503 values to -0.069 (Suppl. Figure 8), which indicates that the quality of all other annotations is at a more comparable standard and, thus, can be used for comparative genomics projects that focus on the Brassicaceae family.

### Interspecific hybridization in *Diplotaxis* and *Moricandida*

It is thought that *D. muralis* and *M. spinosa* are derived from past hybridization events between *D. tenuifolia* and *D. viminea* (Ueno et al., 2006) and between *M. suffruticosa* and *M. nitens* (Perfectti et al., 2017), respectively. The support for tetraploidy by nQuire in both species (Suppl. Figure 1) and their placement close to the respective parental species in the phylogenetic tree (Figure 5) supports these ideas. Further support for tetraploidy in *D. muralis* was found in its large estimated genome size (Suppl. table 4), high synteny with both parental species (Figure 4) and by the large number of HOGs containing only *D. muralis* and either of the parent species (Suppl. Figure 6). Therewith, our work constitutes the first genomic support for the hybrid hypothesis in *D. muralis*, whereas previous support came from isoenzyme pattern and random amplified polymorphic DNA (Eschmann-Grupe et al., 2004). However, the same conclusions cannot be clearly applied to *M. spinosa*. Namely, the smaller genome size estimation and the very high heterozygosity (Suppl. table 4) lead us to speculate that it could be an autopolyploid closely related but not derived from *M. nitens* and *M. suffruticosa* hybridization. It is worth considering that sequence divergence between *M. suffruticosa* and *M. nitens* might be much smaller than between *D. tenuifolia* and *D. viminea*, in which case a hybrid would look more like an autopolyploid. However, *M. spinosa* shared only a moderate amount of HOGs in exclusivity with *M. nitens*, and a small amount with *M. suffruticosa* (Suppl. Figure 6). Therefore, we recommend the estimation of sequence identity between the genomes and divergence time with molecular clocks.

### Evolution of C3-C4 intermediate photosynthesis in the Brassiceae species

Resolving phylogenetic relationships between species is fundamental to evolutionary analysis, which provides a framework to explore the evolution of traits across species. Therefore, we estimated a phylogenetic tree using a multi-species coalescent-based approach (Rannala et al., 2020) to understand how the C3-C4 intermediate photosynthetic trait evolved in the Brassiceae tribe (Figure 5). The relative placement of *H. incana, R. sativus*, and *S. alba* observed in our study agrees with the literature (Huang et al., 2016), whereas the placement of *B. tournefortii* within this clade has not been described earlier. More interestingly, the phylogenetic tree indicates that the C3-C4 intermediate photosynthesis may have evolved independently up to five times in the Brassiceae tribe. Since the Brassiceae don’t contain bona fide C4 species (Sage et al., 2011), we consider it unlikely that the C3-C4 intermediate trait in this tribe has evolved through the hybridization of a C3 and a C4 species (Kadereit et al., 2017). However, we cannot exclude that Brassiceae at some point in time contained C4 species that went extinct. The HIR3 and HIR1 accessions of *H. incana* were placed as sister species in the phylogenetic tree (Figure 5). The time of divergence of these genotypes remains open, but to our knowledge, it could represent the closest pair of diploid C3 and C3-C4 intermediate species known until now in this tribe.

### Further directions in researching C3-C4 photossynthesis in the Brassiceae tribe

Recently, a different accession of *H. incana* was compared to and estimated to have diverged from *B. nigra* 10.35 million years ago (Garassino et al., 2022). However, the close pair HIR3 and HIR1 or even *H. incana* HIR3 with *B. tournefortii* would be even better targeted for genomic comparison, as being phylogenetically closer (Figure 5). Estimating the time of divergence and genomic identity between the two *Hirschfeldia* genotypes of this study would be of great interest for C3-C4 photosynthesis research.

Most earlier studies on comparing C3-C4 photosynthesis within the Brassiceae have focused on the *Moricandia* and *Diplotaxis* genera (Adwy et al., 2015; Razmjoo et al., 1996; Schlüter et al., 2017; Ueno et al., 2006). Both belong to a separate subclade where the species with highest commercial value is rocket salad *E. sativa* (Figure 5). However, the phylogenetic tree of our study indicates the existence of two C3-C4 intermediate species, namely *D. erucoides* and *H. incana* HIR3, in the same subclade where the commercially much more important species of the *Brassica* and *Raphanus* genera exist (Figure 5). Both these species exhibit relatively low carbon compensation points (Schlüter et al. submited), with close relatives like *H*.*incana* HIR1, B. *tournefortii*, or the *Raphanus* available for genomic comparison. Understanding the genetic mechanisms at work in these taxa might be more useful for the engineering of photosynthetic efficiency in the *Brassica* and *Raphanus* crops. Finally, while pairwise comparisons between close C3 and C3-C4 relatives can yield fruitful insights, a more holistic approach comparing multiple taxa and considering their phylogenetic distance is key for statistical power and novel findings. As such, the whole panel of species shows potential in advancing knowledge in C3-C4 evolution.

## CONCLUSION

We generated draft *de novo* genome assemblies using linked- and long-read sequencing data for 18 taxa of the Brassiceae tribe, doubling the sampling depth of genomes within this tribe.Our gene annotation generated high quality models as well as potential to explore variants in genes and regulatory sequences, while our phylogenetic tree indicates that intermediate C3-C4 photosynthesis evolved five times independently across these taxa. This work constitutes the first genomic evidence that *D. muralis* is a hybrid of *D. tenuifolia* and *D. vimin*ea, and it highlights the relevance of *H. incana* HIR3 when discussing C3-C4 photossynthesis. Altogether, the high-quality *de novo* genome assemblies and the gene annotation will be helpful to the scientific community in exploring further the evolution of C3-C4 intermediate photosynthesis in the Brassiceae tribe.

## ACKNOWLEDGEMENTS

The authors give thanks to the Millenium Seed Bank Kew Gardens and the federal ex situ gene bank Getersleben for proving seeds of the species and taxa used in this study. Computational infrastructure and support were provided by the Centre for Information and Media Technology at Heinrich Heine University Düsseldorf. Sequencing support was provided by the Genomics & Transcriptomics Laboratory (GTL) of the Heinrich Heine University Düsseldorf as part of the West German Genome Center (WGGC) and by Max Planck-Genome-Centre Cologne (MP-GC). We thank our colleagues Stephanie Krey and Anja Kyriacidis for their excellent technical assistance. This research was funded by the Deutsche Forschungsgemeinschaft (DFG, German Research Foundation) in the frame of the ERA-CAPS project C4BREED, Germany’s Excellence Strategy (EXC 2048/1, Project ID: 390686111), and a Collaborative Research Centre/Transregio (TRR 341, Project ID: 456082119).

## AUTHOR CONTRIBUTIONS

B.S. and A.P.M.W designed and coordinated the project; U.S. and A.W. and S.T. provided counseling, genetic material, and carbon compensation point sampling.; R.G. performed the genome assemblies and all bioinformatical analyses with inputs from V.S.B. RG and V.S.B wrote the manuscript with contributions from all authors. All authors read and approved the last version of the manuscript.

## DATA AVAILABILITY

The raw SRA data used in this study is deposited in NCBI under BioProject PRJNA905373. The scripts used for genome assembly, annotation and orthology identification were uploaded to github (https://github.com/ViriatoII/C4Evol). The final assemblies have been uploaded to Figshare (https://figshare.com/articles/dataset/C4Evol_Brassicaceae_genomes/21671201).

