## Supplementary Figures for "A genomic panel for studying C3-C4 intermediate photosynthesis in the Brassiceae tribe"

### Slide 1
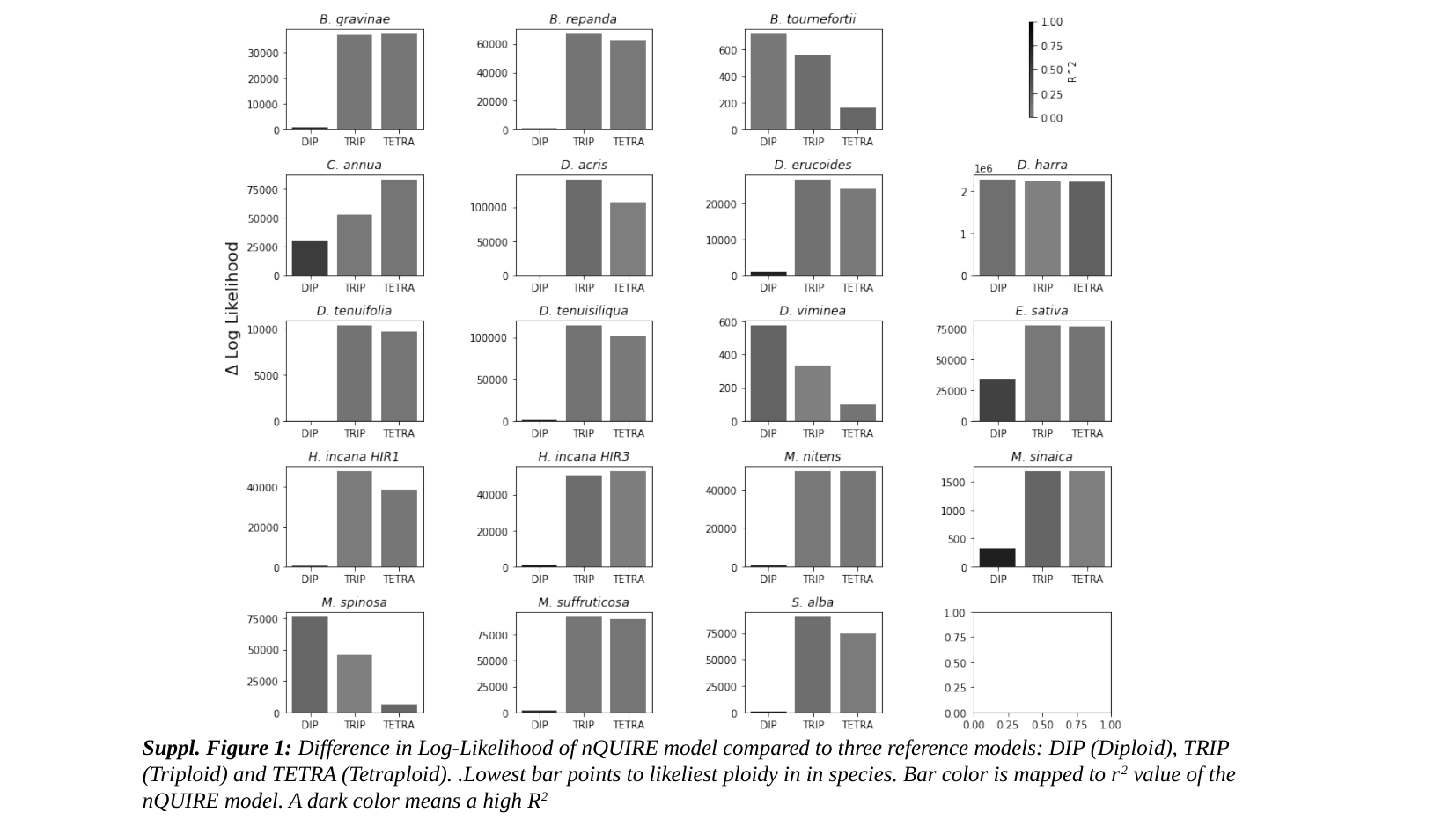

Suppl. Figure 1: Difference in Log-Likelihood of nQUIRE model compared to three reference models: DIP (Diploid), TRIP (Triploid) and TETRA (Tetraploid). .Lowest bar points to likeliest ploidy in in species. Bar color is mapped to r^2^ value of the nQUIRE model. A dark color means a high R^2^

### Slide 2
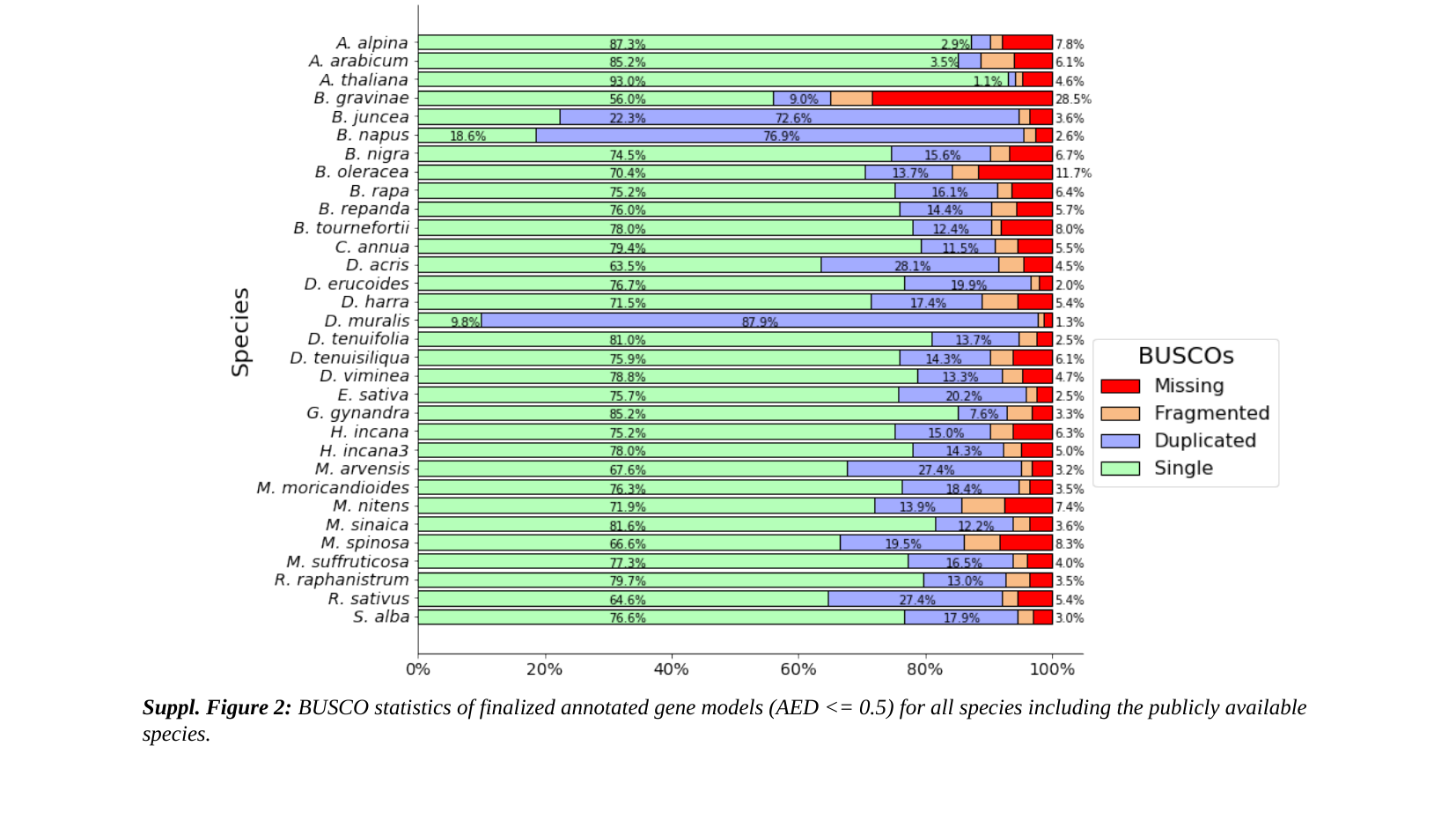

Suppl. Figure 2: BUSCO statistics of finalized annotated gene models (AED <= 0.5) for all species including the publicly available species.

### Slide 3
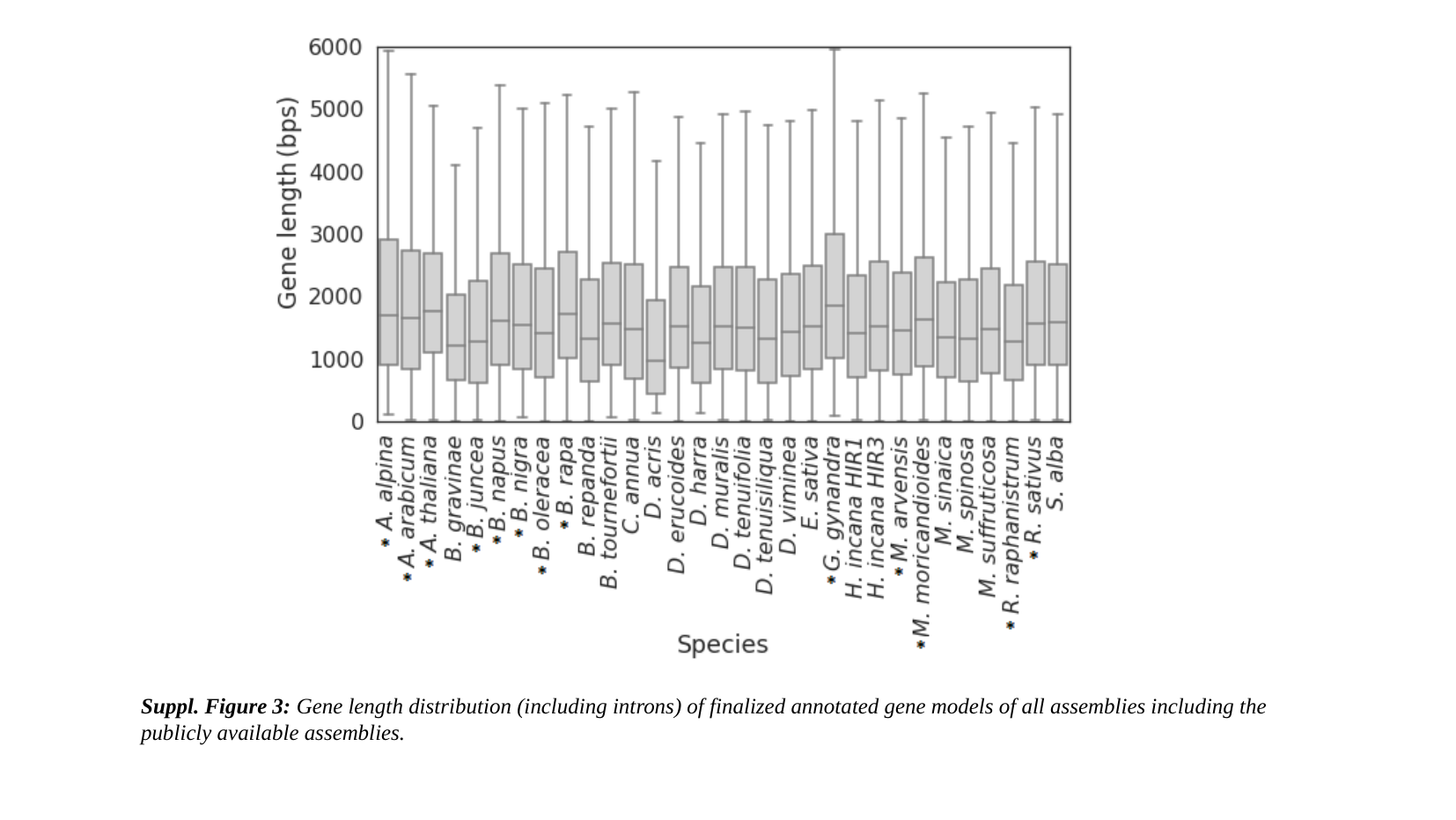

Suppl. Figure 3: Gene length distribution (including introns) of finalized annotated gene models of all assemblies including the publicly available assemblies.

### Slide 4
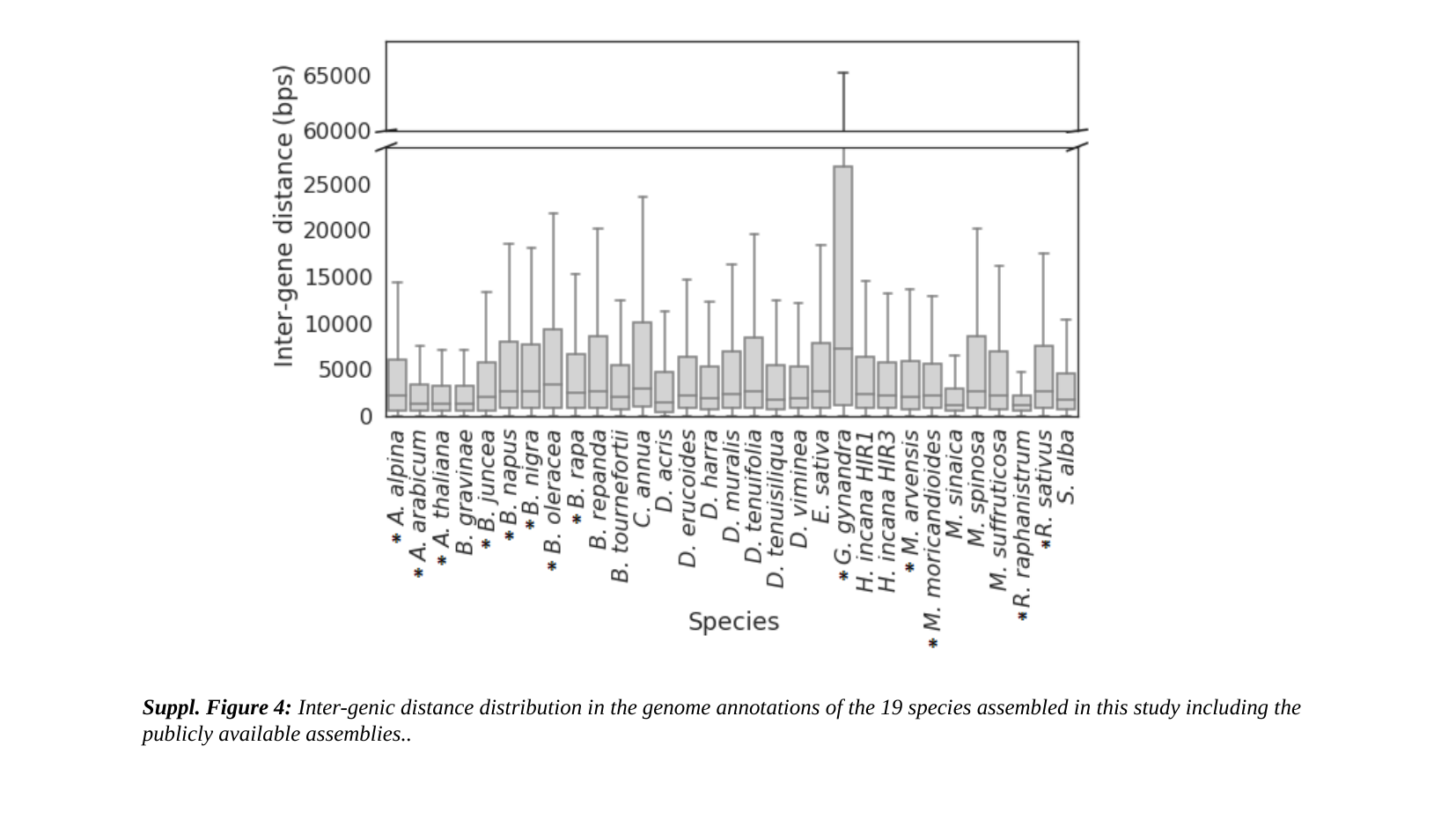

Suppl. Figure 4: Inter-genic distance distribution in the genome annotations of the 19 species assembled in this study including the publicly available assemblies..

### Slide 5
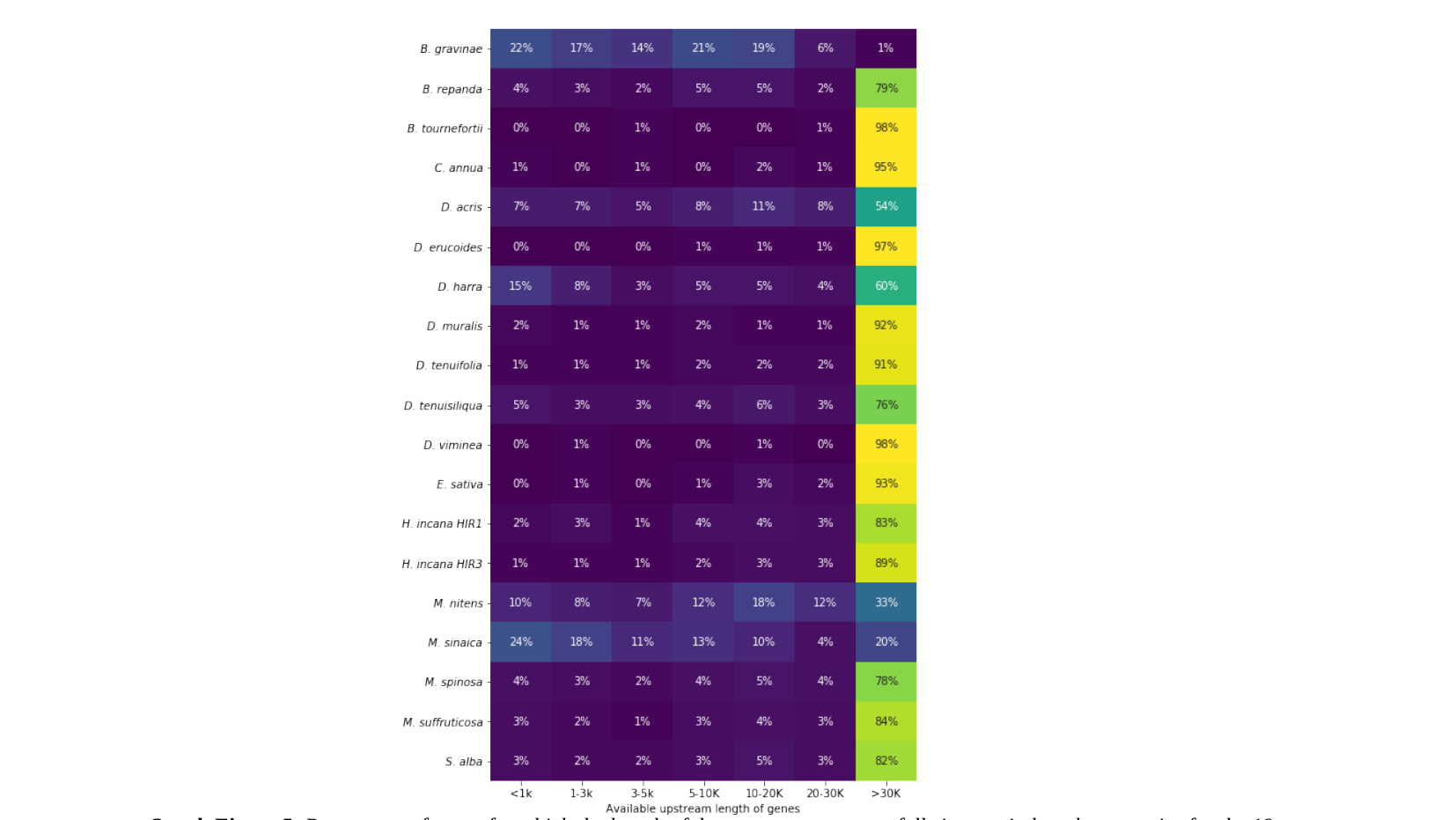

Suppl. Figure 5: Percentage of genes for which the length of the upstream sequence falls in certain length categories for the 19 species assembled in this study.

### Slide 6
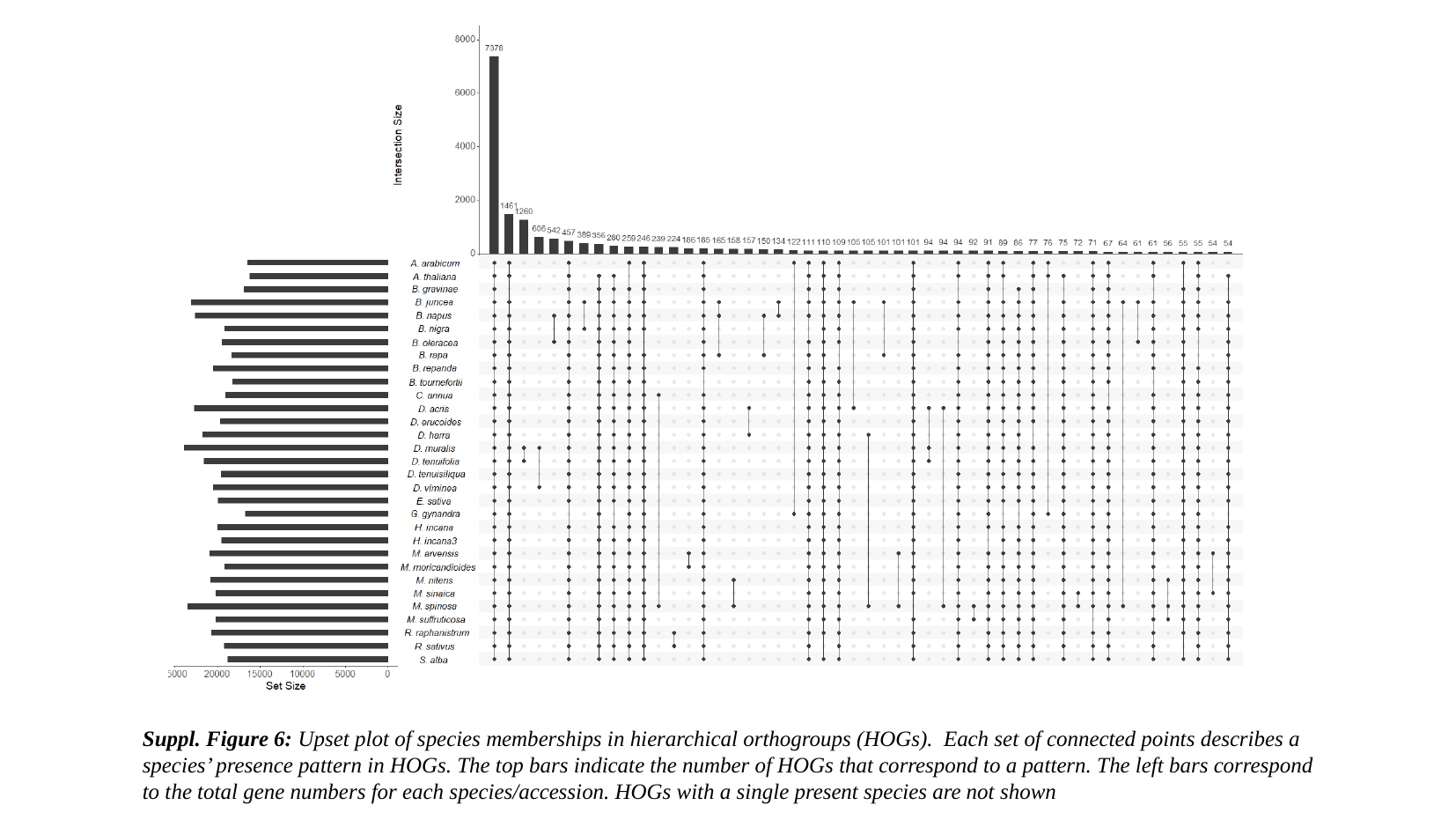

Suppl. Figure 6: Upset plot of species memberships in hierarchical orthogroups (HOGs). Each set of connected points describes a species’ presence pattern in HOGs. The top bars indicate the number of HOGs that correspond to a pattern. The left bars correspond to the total gene numbers for each species/accession. HOGs with a single present species are not shown

### Slide 7
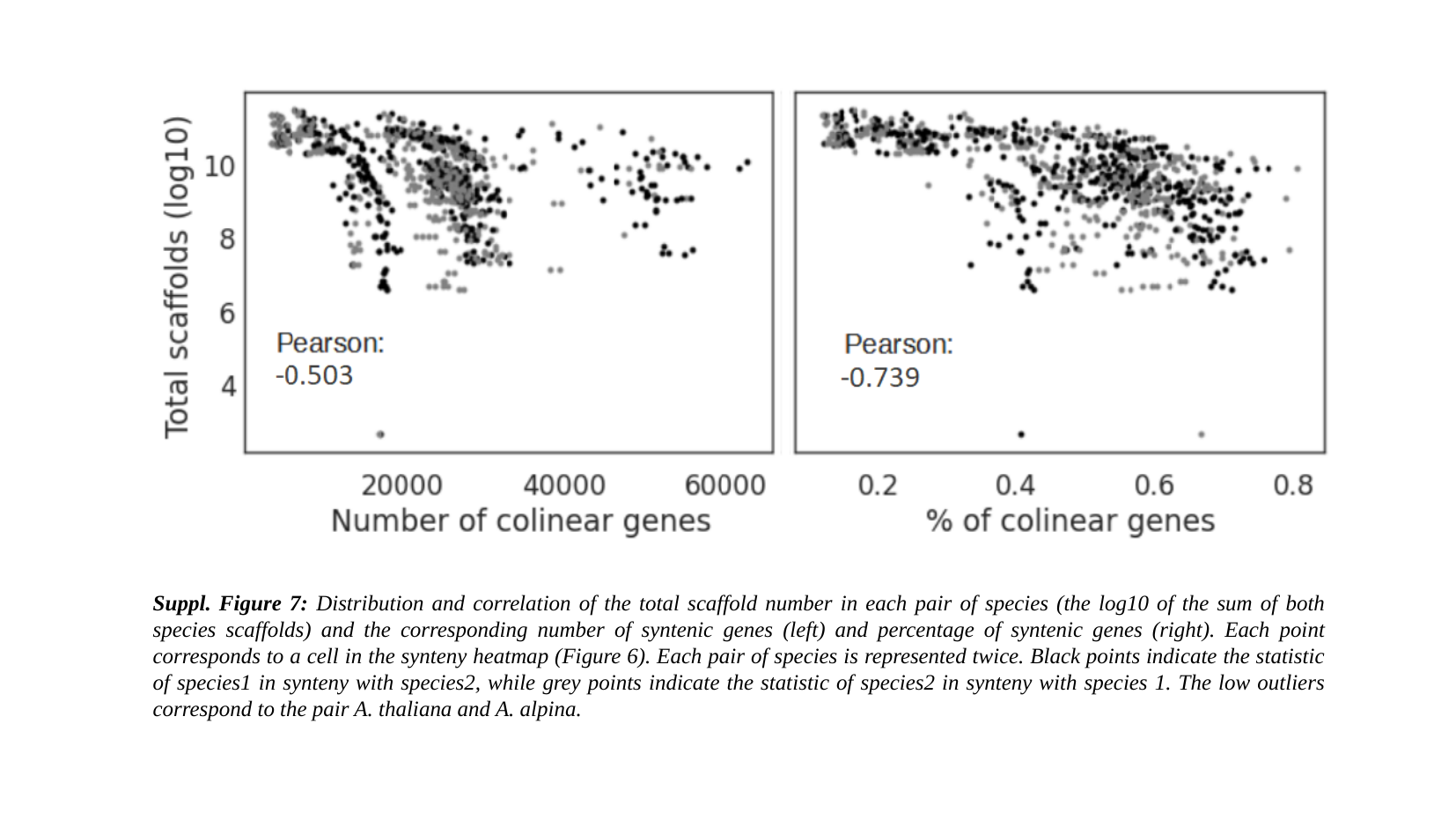

Suppl. Figure 7: Distribution and correlation of the total scaffold number in each pair of species (the log10 of the sum of both species scaffolds) and the corresponding number of syntenic genes (left) and percentage of syntenic genes (right). Each point corresponds to a cell in the synteny heatmap (Figure 6). Each pair of species is represented twice. Black points indicate the statistic of species1 in synteny with species2, while grey points indicate the statistic of species2 in synteny with species 1. The low outliers correspond to the pair A. thaliana and A. alpina.

### Slide 8
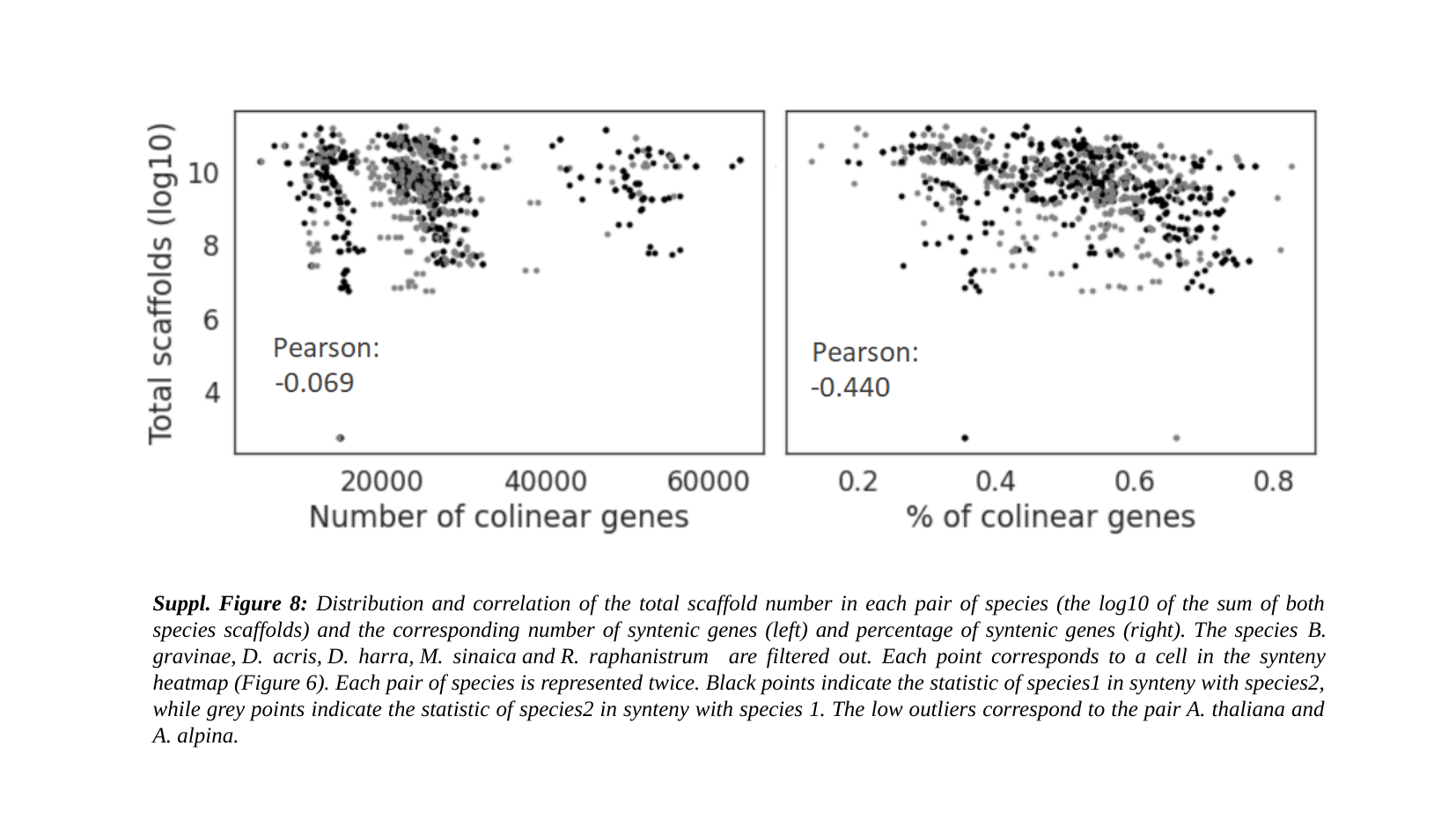

Suppl. Figure 8: Distribution and correlation of the total scaffold number in each pair of species (the log10 of the sum of both species scaffolds) and the corresponding number of syntenic genes (left) and percentage of syntenic genes (right). The species B. gravinae, D. acris, D. harra, M. sinaica and R. raphanistrum are filtered out. Each point corresponds to a cell in the synteny heatmap (Figure 6). Each pair of species is represented twice. Black points indicate the statistic of species1 in synteny with species2, while grey points indicate the statistic of species2 in synteny with species 1. The low outliers correspond to the pair A. thaliana and A. alpina.
